# Synthesis of Novel *N*-Substituted *β*-Amino Acid Derivatives Bearing 2-Hydroxyphenyl Moieties as Promising Antimicrobial Candidates Targeting Multidrug-Resistant Gram-Positive Pathogens

**DOI:** 10.1101/2024.09.25.615057

**Authors:** Povilas Kavaliauskas, Birutė Grybaitė, Birute Sapijanskaite-Banevič, Rūta Petraitienė, Ramunė Grigalevičiūtė, Andrew Garcia, Ethan Naing, Vytautas Mickevičius, Vidmantas Petraitis

## Abstract

The increasing prevalence of antimicrobial resistance among ESKAPE group pathogens presents a significant challenge in the healthcare sector, contributing to higher morbidity and mortality rates globally. Thus, it is essential to develop novel antimicrobial agents effective against drug-resistant pathogens. In this study, we report the synthesis and *in vitro* antimicrobial activity characterization of novel *N*-substituted *β*-amino acid derivatives bearing 2-hydroxyphenyl core against multidrug-resistant bacterial pathogens. The synthesized compounds (**2–26**) exhibited promising antimicrobial activity specifically against Gram-positive bacteria, with minimum inhibitory concentrations (MIC) ranging from 4 to 128 µg/mL. None of the compounds demonstrated activity against Gram-negative pathogens or drug-resistant fungi. Compounds **9** (R = 4-nitrophenyl), **17** (R = 5-nitro-2-thienyl), **18** (R = 5-nitro-2-furyl), thiosemicarbazide **16**, and **26** exhibited the most promising activity against *Staphylococcus aureus* MRSA USA300 lineage strain TCH-1516, with MIC values between 4 and 16 µg/mL. Compound **26** demonstrated strong antimicrobial activity against both *S. aureus* TCH-1516 and *E. faecalis* AR-0781, with the activity comparable to control antibiotics. These findings indicate that *N*-substituted *β*-amino acid derivatives with a 2-hydroxyphenyl core warrant further investigation as a potential scaffold for the further development of antimicrobial agents based on compound **26** targeting Gram-positive pathogens.

## Introduction

The growing prevalence of antimicrobial resistance represents a critical challenge in healthcare setting, with the increasing numbers of infections caused by ESKAPE group pathogens [1–3]. This group comprises of drug-resistant *Enterococcus* species, *Staphylococcus aureus*, *Klebsiella pneumoniae*, *Acinetobacter baumannii*, *Pseudomonas aeruginosa*, and *Enterobacter* species [4,5]. These pathogens are especially challenging for their antimicrobial resistance to multiple first line antibiotics, thereby complicating treatment regimens and leading to higher rates of morbidity and mortality [5]. The clinical impact of ESKAPE pathogens is profound, with methicillin-resistant *Staphylococcus aureus* (MRSA) and vancomycin-resistant *Enterococcus* (VRE) being prominent contributors to severe healthcare-associated infections [4, 6–7]. The high mortality rates associated with these infections, coupled with the prolonged hospital stays and increased healthcare costs, underscore the urgent need for the development of novel antimicrobial agents for further pre-clinical development and optimization [8,9]. Infections caused by MRSA are known to cause a variety of infections, including skin and soft tissue infections, pneumonia, endocarditis, osteomyelitis, and sepsis, with mortality rates ranging from 20% to 50% depending on the severity and site of infection [10–12]. VRE predominantly causes bloodstream infections, urinary tract infections, intra-abdominal infections, and wound infections, with mortality rates for VRE bacteremia reaching up to 60% [13–15].

The molecular mechanisms by which these pathogens evade antibiotics are complex and multifaceted [16]. MRSA confers resistance to *β*-lactam antibiotics primarily through the acquisition of the *mecA* gene, which encodes a penicillin-binding protein (PBP2a) with low affinity for *β*-lactams, thereby preventing the inhibition of cell wall synthesis [16,17]. Additionally, MRSA employs various other resistance mechanisms, such as efflux pumps and the production of various antibiotics-modifying enzymes, often conferring resistance to aminoglycosides [18,19]. VRE, on the other hand, utilizes modification in cell wall precursors biosynthesis to evade vancomycin-mediated killing [20]. The acquisition of *vanA* or *vanB* gene clusters results in the modification of the terminal D-alanine-D-alanine to D-alanine-D-lactate in the peptidoglycan precursor, significantly reducing vancomycin binding affinity in VRE cell wall biogenesis pathway [20–22]. This modification impedes the antibiotic’s ability to inhibit cell wall synthesis, thereby conferring resistance to vancomycin [20]. The clinical impact of these resistance mechanisms is profound, as they severely limit the therapeutic options available for treating MRSA and VRE infections, necessitating the development of new antimicrobial strategies [21,22].

Various amino acid derivatives have been extensively investigated in medicinal chemistry as novel pharmacophores for targeting a variety of diseases, including cancers, bacterial and fungal infections, and parasitic diseases [22, 24–27]. These derivatives are attractive candidates for drug design due to their inherent biological activity, structural diversity, and ability to mimic natural substrates in biological systems [24, 26–27]. For instance, modifications of amino acid side chains or addition of various aromatic or heterocyclic substituents can enhance binding affinity to specific biological targets, improve selectivity, and increase metabolic stability [22, 24, 27]. Among substituted amino acid derivatives, *N*-substitution in *β*-amino acids offers promising scaffold with remarkable versatility in tailoring molecular properties. This *N*-substituted scaffold allows to incorporate a vide variety of substitutions that are crucial for further optimization of physicochemical attributes, influencing lipophilicity, electronic distribution, and overall bioactivity.

Our previous studies have demonstrated that amino acid derivatives bearing a 3-hydroxyphenyl group exhibit promising antimicrobial activity against multidrug-resistant bacterial pathogens [28]. Additionally, the synthesized compounds exhibited significant antifungal activity against highly multidrug-resistant fungal pathogens, including the emerging azole-resistant *Candida auris* [28]. Further in silico analyses indicated favorable predicted pharmacological and drug-like properties, establishing 3-((4-hydroxyphenyl)amino)propanoic acid as a promising scaffold for the development of antimicrobial candidates [28]. Building on these findings, the present study explores the synthesis pathways and *in vitro* antimicrobial properties of *N*-substituted *β*-amino acid derivatives bearing a 2-hydroxyphenyl moiety, along with various aromatic and heterocyclic substituents. In this study, we successfully demonstrate that *N*-substituted *β*-amino acid derivatives with a 2-hydroxyphenyl core, as well as heterocyclic substituents, could serve as promising scaffolds for the development of novel antimicrobial candidates targeting Gram-positive pathogens.

## Materials and methods

### Chemical synthesis

#### General analytical procedures

The reaction course and purity of the synthesized compounds were monitored by TLC using aluminum plates precoated with Silica gel with F254 nm (Merck KGaA, Darmstadt, Germany). Reagents and solvents were obtained from Sigma-Aldrich (St. Louis, MO, USA) and used without further purification. Melting points were determined with a B-540 melting point analyzer (Büchi Corporation, New Castle, DE, USA) and were uncorrected. IR spectra (υ, cm^−1^) were recorded on a Perkin–Elmer Spectrum BX FT–IR spectrometer using KBr pellets. NMR spectra were recorded on a Brucker Avance III (400, 101 MHz) spectrometer. Chemical shifts were reported in (δ) ppm relative to tetramethylsilane (TMS) with the residual solvent as internal reference ([D_6_]DMSO, δ = 2.50 ppm for ^1^H and δ = 39.5 ppm for ^13^C). Data were reported as follows: chemical shift, multiplicity, coupling constant (Hz), integration, and assignment. Elemental analyses (C, H, N) were conducted using the Elemental Analyzer CE-440; their results were found to be in good agreement (±0.3%) with the calculated values.

#### General procedures for preparation of compounds 2–26

##### 3,3’-((2-Hydroxyphenyl)azanediyl)dipropionic acid (**2**)

A mixture of *o*-aminophenol (1) (10.9 g, 100 mmol), acrylic acid (18 g, 250 mmol) and water (100 mL) was heated under reflux for 14 h and cooled. Crystaline product 2 was filtered off, washed with propan-2-ol, and dried. White powder, yield 18.97 g (75%), m.p. 181−183 °C (from propan-2-ol). ^1^H NMR (400 MHz, DMSO-*d_6_*) δ: 2.29 (t, *J* = 7.1 Hz, 4H, 2x CH_2_CO), 3.18 (t, *J* = 7.1 Hz, 4H, 2x NCH_2_), 6.66–7.06 (m, 4H, H_Ar_); 8.77 (br.s, 1H, OH); 12.06 (br.s, 2H, 2x OH); ^13^C NMR (101 MHz, DMSO-*d_6_*) δ: 32.39 (<u>C</u>H_2_CO), 48.63 (NCH_2_), 115.60, 119.36, 123.13, 124.94, 136.50, 152.73 (C_Ar_), 173.77 (2x C=O). IR (KBr): ν_max_ (cm^−1^) = 3045, 2971 (3x OH), 1698 (2x C=O). Anal. Calcd. for C_12_H_15_NO_5_, %: C 56.91; H 5.97; N 5.53. Found: C 56.69; H 5.70; N 5.33.

##### 3-(2-Oxo-3,4-dihydrobenzo[b][1,4]oxazepin-5(2H)-yl)-N-(4-sulfamoylphenyl)propenamide (**3**)

The dipropionic acid 2 (0.5g 2, mmol) is dissolved in dimethylformamide (2 mL) by heating, then sulfanilamide (0.83g, 4.8 mmol,) is dissolved in dimethylformamide (2 mL) in another beaker and both prepared solutions are poured into a flask and triethylamine (12 mmol) is added dropwise, the reaction mixture 20 minutes stirred at room temperature. Separately, HBTU (2.28 g, 6 mmol) was dissolved in 9 mL of dimethylformamide and added dropwise to the reaction mixture. The reaction was stirred at room temperature for 24 h. After that, the reaction mixture is diluted with water (40 mL), the formed crystals are filtered and washed with 5% potassium carbonate solution, water and dried. White powder, yield 0.22 g (28%), m.p. 190−192 °C (from dioxane/water mixture). ^1^H NMR (400 MHz, DMSO-*d_6_*) δ: 2.58 (t, *J* = 7.0 Hz, 4H, 2x CH_2_CO), 3.39 (t, *J* = 7.0 Hz, 4H, 2x NCH_2_), 6.99–7.16 (m, 2H, SO_2_NH_2_), 7.19–7.32 (m, 4H, H_Ar_), 7.72 (q, *J* = 8.7 Hz, 4H, H_Ar_), 12.21 (s, 1H, NH); ^13^C NMR (101 MHz, DMSO-*d_6_*) δ: 31.67 (<u>C</u>H_2_CONH), 35.02 (CH_2_<u>C</u>ONHO), 47.46 (NCH_2_ CH_2_CONH), 54.29 (N<u>C</u>H_2_CH_2_CONHO), 118.63, 119.45, 121.13, 123.91, 126.29, 126.64, 138.20, 138.42, 141.96, 146.86 (C_Ar_), 169.94, 170.02 (2x C=O). IR (KBr): ν_max_ (cm^−1^) = 3344 (NH_2_), 3241 (NH), 1742, 1689 (2x C=O). Anal. Calcd. for C_18_H_19_N_3_O_5_S, %: C 55.52; H 4.92; N 10.79. Found: C 55.31; H 4.72; N 10.55. HRMS m/z calculated for C_18_H_19_N_3_O_5_S [M+H]^+^: 390.1045; found: 390.1119.

##### Dimethyl 3,3’-((2-hydroxyphenyl)azanediyl)dipropionate (**4**)

A mixture of dipropionic acid 2 (17.48 g, 69 mmol), conc. sulfuric acid (8.6 g, 4.7 mL, 87 mmol) and methanol (250 mL) was heated under reflux for 7 h. Then the solvent was evaporated under reduced pressure, and the residuse neutralized with 5% sodium carbonate solution to pH 7. The obtained solid was filtered off, washed with plenty of water and recrystallized from propan-2-ol. Light brown powder, yield 15.7 g (81%), m.p. 87–89 °C. ^1^H NMR (400 MHz, DMSO-*d_6_*) δ: 2.37 (t, *J* = 7.0 Hz, 4H, 2x CH_2_CO), 3.22 (t, *J* = 7.0 Hz, 4H, 2x NCH_2_), 3.54 (s, 6H, 2x CH_3_), 6.68–7.05 (m, 4H, H_Ar_); 8.66 (s, 1H, OH); ^13^C NMR (101 MHz, DMSO-*d_6_*) δ: 32.89 (<u>C</u>H_2_CO), 48.41 (NCH_2_), 51.26 (CH_3_), 115.53, 119.17, 123.27, 124.75, 135.94, 152.56 (C_Ar_), 172.35 (2x C=O). IR (KBr): ν_max_ (cm^−1^) = 3308 (OH), 1727 (2x C=O). Anal. Calcd. for C_14_H_19_NO_5_, %: C 59.78; H 6.81; N 4.98. Found: C 59.52; H 6.60; N 4.72.

##### 3,3’-((2-Hydroxyphenyl)azanediyl)di(propanehydrazide) (**5**)

A mixture of methyl ester 4 (7.28 g, 26 mmol), hydrazine hydrate (7.89 g, 157 mmol), and propan-2-ol (25 mL) was heated under reflux for 5 h and cooled. Crystalline product 5 was filtered off, washed with propan-2-ol, diethyl ether, and dried. White powder, yield 5.82 g (80%), m.p. 149−151 °C (from propan-2-ol). ^1^H NMR (400 MHz, DMSO-*d_6_*) δ: 2.13 (t, *J* = 7.1 Hz, 4H, 2x CH_2_CO), 3.12 (t, *J* = 7.2 Hz, 4H, 2x NCH_2_), 4.20 (s, 4H, 2x NH_2_), 6.70–7.08 (m, 4H, H_Ar_); 9.01 (s, 3H, OH, 2x N*H*NH_2_); ^13^C NMR (101 MHz, DMSO-*d_6_*) δ: 31.57 (<u>C</u>H_2_CO), 49.00 (NCH_2_), 115.69, 119.12, 122.64, 124.64, 136.65, 152.83 (C_Ar_), 170.62 (2x C=O). IR (KBr): ν_max_ (cm^−1^) = 3295 (OH), 3178 (NH_2_), 3057 (NH_2_), 1677, 1624 (C=O). Anal. Calcd. for C_12_H_19_N_5_O_3_, %: C 51.23; H 6.81; N 24.90. Found: C 51.01; H 6.62; N 24.69.

##### 2-((2-(1H-benzo[d]imidazol-2-yl)ethyl)amino)phenol (**6**)

Solution of compound 2 (0.7 g, 0.0028 mol) and *o*-phenylenediamine (0.61 g, 0.0056 mol) in dilute hydrochloric acid (1:1, 8 mL) was heated at reflux for 24 h, then cooled down. The reaction mixture is neutralized with a 5% sodium carbonate solution to pH 8. The formed crystals are recrystallized from a mixture of 1,4-dioxane and water (1:1, 15 ml). Brown powder, yield 0.2 g (28%), m.p. 196−198 °C. ^1^H NMR (400 MHz, DMSO-*d_6_*) δ: 3.10 (t, *J* = 7.0 Hz, 2H, 2x CH_2_CN), 3.52 (q, *J* = 7.0 Hz, 2H, 2x NCH_2_), 4.86 (t, *J* = 7.0 Hz, 1H, NHCH_2_), 6.23–6.74 (m, 4H, H_Ar_); 7.01–7.22 (m, 2H, H_Ar_), 7.29–7.68 (m, 2H, H_Ar_), 9.23 (s, 1H, OH), 12.28 (s, 1H, NH);^13^C NMR (101 MHz, DMSO-*d_6_*) δ: 28.52 (<u>C</u>H_2_CO), 41.34 (NCH_2_), 109.82, 110.81, 113.50, 116.02, 118.14, 119.75, 120.91, 121.53, 134.23, 137.09, 143.33, 144.22, 153.44 (C_Ar_). IR (KBr): ν_max_ (cm^−1^) = 3310 (OH), 3055 (2x NH). Anal. Calcd. for C_15_H_15_N_3_O, %: C 71.13; H 5.97; N 16.59. Found: C 70.95; H 5.70; N 16.32.

#### General procedure for the preparation of hydrazones **7–22**

To a solution of hydrazide 5 (0.42g, 1.5 mmol) in 2-propanol (15 mL), the corresponding aromatic aldehyde was added (1.65 mmol) and the mixture was heated at reflux for 2 h, then cooled down, and the formed precipitate was filtered off, washed with methanol, diethyl ether and recrystallizing from 1,4-dioxane.

##### 3,3’-((2-Hydroxyphenyl)azanediyl)bis(N’-(benzylidene)propanehydrazide) (**7**)

White powder, yield 0.57 g (83%), m.p. 223−225 °C. ^1^H NMR (400 MHz, DMSO-*d_6_*) δ: 2.34 and 2.77 (2t, *J* = 7.1 Hz, 2H, 2x CH_2_CO), 3.17–3.41 (m, 4H, 2x NCH_2_), 6.68–7.21 (m, 4H, H_Ar_); 7.29–7.79 (m, 10H, H_Ar_), 7.83–8.19 (m, 2H, 2x CH), 8.91, 8.96, 9.02 (3s, 1H, OH), 11.30, 11.31, 11.41 (3s, 2H, 2x NH); ^13^C NMR (101 MHz, DMSO-*d_6_*) δ: 30.19, 30.42, 31.59, 32.35, 32.55 (<u>C</u>H_2_CO), 48.27, 48.49, 48.67, 48.92 (NCH_2_), 115.62, 119.07, 119.18, 122.44, 122.75, 124.33, 124.55, 126.64, 126.68, 126.99, 128.74, 128.77, 129.67, 129.90, 134.18, 134.32, 136.21, 136.50, 136.90, 152.84, 145.97, 152.37, 152.54, 152.79, 167.52, 167.62 (C_Ar_) 170.60, 173.44, 173.49 (C=O). IR (KBr): ν_max_ (cm^−1^) = 3109 (OH), 3007 (2x NH), 1672 (2x C=O). Anal. Calcd. for C_26_H_27_N_5_O_3_, %: C 68.25; H 5.95; N 15.31. Found: C 68.00; H 5.73; N 15.11. HRMS m/z calculated for C_26_H_27_N_5_O_3_ [M+H]^+^: 458.2114; found: 458.2184.

##### 3,3’-((2-Hydroxyphenyl)azanediyl)bis(N’-(2,4-difluorobenzylidene)propanehydrazide) (**8**)

White powder, yield 0.57 g (83%), m.p. 224−226 °C. ^1^H NMR (400 MHz, DMSO-*d_6_*) δ: 2.23–2.39 and 2.64–2.87 (2m, 4H, 2x CH_2_CO), 3.15–3.34 (m, 4H, 2x NCH_2_), 6.68–7.37 (m, 8H, H_Ar_); 7.58–7.95 (m, 2H, H_Ar_), 7.99–8.11 and 8.20–8.30 (2m, 2H, 2x N=CH), 8.88, 8.90, 9.00 (3s, 1H, OH), 11.39, 11.41, 11.53 (3s, 2H, 2x NH); ^13^C NMR (101 MHz, DMSO-*d_6_*) δ: 30.14, 30.37, 32.44, 32.62 (<u>C</u>H_2_CO), 48.16, 48.43, 48.63, 48.76 (NCH_2_), 104.12, 104.36, 104.63, 112.54, 115.57, 118.54, 118.64, 119.07, 119.21, 122.31, 122.70, 124.39, 124.59, 127.48, 127.84, 127.88, 134.83, 136.49, 136.82, 137.95, 152.35, 152.51, 159.33, 159.46, 161.64, 161.97, 164.12, 167.64, 167.69 (C_Ar_), 173.53, 173.58 (2x C=O). IR (KBr): ν_max_ (cm^−1^) = 3297 (OH), 3196 (2x NH), 1673 (2x C=O). Anal. Calcd. for C_26_H_23_F_4_N_5_O_3_, %: C 58.98; H 4.38; N 13.23. Found: C 58.73; H 4.13; N 13.03. HRMS m/z calculated for C_26_H_23_F_4_N_5_O_3_ [M+H]^+^: 530.1737; found: 530.1810.

##### 3,3’-((2-Hydroxyphenyl)azanediyl)bis(N’-(4-nitrobenzylidene)propanehydrazide) (**9**)

Light yellow powder, yield 0.55 g (67%), m.p. 225−227 °C (from propan-2-ol). ^1^H NMR (400 MHz, DMSO-*d_6_*) δ: 2.38 (t, *J* = 6.8 Hz, 2H, CH_2_CO), 2.80 (t, *J* = 6.7 Hz, 2H, CH_2_CO), 3.25−3.39 (m, 4H, 2x NCH_2_), 6.73–8.30 (m, 14H, H_Ar_, 2x N=CH); 8.90, 8.93, 9.06 (3s, 1H, OH), 11.58, 11.60, 11.71 (3s, 2H, 2x NH); ^13^C NMR (101 MHz, DMSO-*d_6_*) δ: 30.25, 30.45, 32.45, 32.63 (<u>C</u>H_2_CO), 48.10, 48.36, 48.66, 48.79 (NCH_2_), 115.58, 115.64, 115.79, 119.10, 119.25, 119.36, 122.29, 122.79, 123.13, 123.88, 123.93, 123.96, 124.33, 124.60, 127.42, 127.51, 127.84, 136.22, 136.46, 136.82, 140.39, 140.46, 140.51, 140.69, 143.47, 143.57, 147.44, 147.50, 147.62, 147.70, 152.29, 152.50, 152.78, 168.02, 168.07 (C_Ar_), 173.83, 173.90 (2x C=O). IR (KBr): ν_max_ (cm^−1^) = 3265 (OH), 3092 (2x NH), 1675 (2x C=O). Anal. Calcd. for C_26_H_25_N_7_O_7_, %: C 57.04; H 4.60; N 17.91. Found: C 56.83; H 4.42; N 17.75.

##### 3,3’-((2-Hydroxyphenyl)azanediyl)bis(N’-(4-chlorobenzylidene)propanehydrazide) (**10**)

White powder, yield 0.55 g (69%), m.p. 220−222 °C (from dioxane). ^1^H NMR (400 MHz, DMSO-*d_6_*) δ: 2.34 (t, *J* = 6.8 Hz, 2H, CH_2_CO), 2.76 (t, *J* = 7.1 Hz, 2H, CH_2_CO), 3.25−3.36 (m, 4H, 2x NCH_2_), 6.70–8.12 (m, 14H, H_Ar_, 2x N=CH); 8.89, 8.94, 9.02 (3s, 1H, OH), 11.35, 11.36, 11.47 (3s, 2H, 2x NH); ^13^C NMR (101 MHz, DMSO-*d_6_*) δ: 30.18, 30.41, 32.36, 32.55 (<u>C</u>H_2_CO), 48.21, 48.43, 48.66, 48.87 (NCH_2_), 115.56, 115.62, 115.74, 119.08, 119.20, 119.31, 122.35, 122.73, 123.10, 124.34, 124.57, 124.86, 128.21, 128.26, 128.59, 128.77, 128.80, 128.83, 133.10, 133.13,133.25, 133.27, 134.06, 134.31, 136.22, 136.50, 136.85, 144.66, 144.77, 152.33, 152.51, 152.77, 167.62, 167.69 (C_Ar_), 173.49, 173.54 (2x C=O). IR (KBr): ν_max_ (cm^−1^) = 3179 (OH), 3107 (2x NH), 1673 (2x C=O). Anal. Calcd. for C_26_H_25_ Cl_2_N_5_O_3_, %: C 59.32; H 4.79; N 13.30. Found: C 59.11; H 4.54; N 13.17. HRMS m/z calculated for C_26_H_25_ Cl_2_N_5_O_3_ [M+Na]^+^: 548.1334; found: 548.1228.

##### 3,3’-((2-Hydroxyphenyl)azanediyl)bis(N’-(4-(dimethylamino)benzylidene)propanehydrazide) (**11**)

White powder, yield 0.75 g (91%), m.p. 206−208 °C. ^1^H NMR (400 MHz, DMSO-*d_6_*) δ: 2.30 and 2.73 (2q, *J* = 6.5 Hz, 2H, 2x CH_2_CO), 2.91, 2.93, 2.95 (3s, 12H, 4x CH_3_); 3.20–3.33 (m, 4H, 2x NCH_2_), 6.56–6.98 (m, 6H, H_Ar_); 7.06–7.17 (m, 1H, H_Ar_), 7.29–7.51 (m, 4H, H_Ar_), 7.76–8.01 (m, 2H, 2x N=CH), 9.00, 9.02, 9.06 (3s, 1H, OH), 11.03, 11.12, 11.13 (3s, 2H, 2xNH); ^13^C NMR (101 MHz, DMSO-*d_6_*) δ: 30.26, 30.50, 32.31, 32.36, 32.44, 32.51 (4x CH_3_, <u>C</u>H_2_CO), 48.24, 48.59, 49.02 (NCH_2_), 111.73, 111.75, 111.77, 115.64, 119.17, 121.60, 121.63, 122.28, 122.58, 123.07, 124.31, 124.51, 127.92, 127.95, 128.31, 128.35, 136.32, 136.72, 137.15, 143.79, 146.85, 147.02, 151.22, 151.38, 152.52, 166.95, 167.03 (C_Ar_), 172.87 (2x C=O). IR (KBr): ν_max_ (cm^−1^) = 3437 (OH), 3078 (2x NH), 1661 (2x C=O). Anal. Calcd. for C_30_H_37_N_7_O_3_, %: C 66.28; H 6.86; N 18.03. Found: C 66.04; H 6.65; N 17.86. HRMS m/z calculated for C_30_H_37_N_7_O_3_ [M+H]^+^: 544.2957; found: 544.3025.

##### 3,3’-((2-Hydroxyphenyl)azanediyl)bis(N’-(4-hydroxybenzylidene)propanehydrazide) (**12**)

White powder, yield 0.45 g (62%), m.p. 164−166 °C (from methanol). ^1^H NMR (400 MHz, DMSO-*d_6_*) δ: 2.31 (t, *J* = 6.9 Hz, 2H, CH_2_CO), 2.74 (t, *J* = 6.7 Hz, 2H, CH_2_CO), 3.19−3.36 (m, 4H, 2x NCH_2_), 6.69–8.07 (m, 14H, H_Ar_, 2x N=CH); 8.98 (t, 1H, OH), 9.84, 9.87 (2s, 2H, 2x OH), 11.10, 11.21 (2s, 2H, 2x NH); ^13^C NMR (101 MHz, DMSO-*d_6_*) δ: 30.21, 30.43, 32.34, 32.53 (<u>C</u>H_2_CO), 48.26, 48.55, 48.69, 49.02 (NCH_2_), 115.56, 115.63, 115.74, 119.06, 119.17, 119.31, 122.38, 122.71, 123.13, 125.24, 125.27, 125.29, 128.37, 128.40, 128.75, 136.30, 136.62, 137.06, 143.24, 146.37, 146.50, 152.40, 152.58, 152.82, 159.05, 159.27, 167.18, 167.29 (C_Ar_), 173.12, 173.15 (2x C=O). IR (KBr): ν_max_ (cm^−1^) = 3577 (OH), 3088 (2x NH), 1654 (2x C=O). Anal. Calcd. for C_26_H_27_N_5_O_5_, %: C 63.79; H 5.56; N 14.31. Found: C 63.55; H 5.36; N 14.15. HRMS m/z calculated for C_26_H_27_N_5_O_5_ [M+Na]^+^: 512.2012; found: 512.1903.

##### 3,3’-((2-Hydroxyphenyl)azanediyl)bis(N’-(3,4,5-trimethoxybenzylidene)propanehydrazide) (**13**)

White powder, yield 0.8 g (83%), m.p. 214−216 °C (from methanol). ^1^H NMR (400 MHz, DMSO-*d_6_*) δ: 2.33 and 2.77 (q, *J* = 6.9 Hz, 4H, CH_2_CO), 3.12−3.36 (m, 4H, 2x NCH_2_), 3.60–3.85 (m, 18H, 6x OCH_3_), 6.66–7.17 (m, 8H, H_Ar_); 7.83, 7.87, 8.02, 8.04 (4s, 2H, 2xN=CH); 8.81, 8.92, 9.02 (3s, 1H, OH), 11.31, 11.34, 11.36, 11.39 (4s, 2H, 2x NH); ^13^C NMR (101 MHz, DMSO-*d_6_*) δ: 30.17, 30.37, 32.39, 32.57 (<u>C</u>H_2_CO), 48.68, 48.88, 55.79, 55.85, 55.88, 55.90, 60.08 (NCH_2_, 6x OCH_3_), 103.72, 103.79, 104.16, 115.27, 115.50, 119.01, 119.16, 122.95, 124.68, 129.74, 129.82, 136.34, 136.46, 136.68, 138.80, 138.85, 139.03, 142.47, 142.57, 145.97, 146.04, 152.73, 153.07, 153.11, 167.43, 167.51 (C_Ar_), 173.35, 173.43 (2x C=O). IR (KBr): ν_max_ (cm^−1^) = 3327 (OH), 3111 (2x NH), 1670 (2x C=O). Anal. Calcd. for C_32_H_39_N_5_O_9_, %: C 60.27; H 6.16; N 10.98. Found: C 60.03; H 5.94; N 10.71.

##### 3,3’-((2-Hydroxyphenyl)azanediyl)bis(N’-(naphthalen-1-ylmethylene)propanehydrazide) (**14**)

White powder, yield 0.73 g (87%), m.p. 213−215 °C (from methanol). ^1^H NMR (400 MHz, DMSO-*d_6_*) δ: 2.42 and 2.86 (t, *J* = 7.2 Hz, 4H, CH_2_CO), 3.34−3.52 (m, 4H, 2x NCH_2_), 6.71–6.99 (m, 3H, H_Ar_); 7.11–7.25 (m, 1H, H_Ar_); 7.44–8.05 (m, 12H, H_Ar_); 8.47–8.85 (m, 4H, H_Ar_, 2x N=CH); 8.94, 9.02, 9.10 (3s, 1H, OH), 11.35, 11.37, 11.53, 11.54 (4s, 2H, 2x NH); ^13^C NMR (101 MHz, DMSO-*d_6_*) δ: 30.34, 30.56, 32.42, 32.61 (<u>C</u>H_2_CO), 48.45, 48.63, 48.81, 48.97 (NCH_2_), 115.51, 115.63, 115.78, 119.15, 119.24, 122.68, 122.90, 123.64, 124.30, 124.54, 124.68, 125.46, 126.14, 126.19, 127.01, 127.11, 127.18, 127.26, 127.86, 128.75, 129.34, 129.36, 129.52, 129.95, 129.97, 130.07, 130.10, 130.36, 133.44, 133.47, 133.50, 136.46, 136.75, 142.62, 142.71, 145.96, 146.08, 152.62, 152.68, 167.54, 167.63 (C_Ar_), 173.32, 173.37 (2x C=O). IR (KBr): ν_max_ (cm^−1^) = 3169 (OH), 3092 (2x NH), 1675 (2x C=O). Anal. Calcd. for C_34_H_31_N_5_O_3_, %: C 73.23; H 5.60; N 12.56. Found: C 73.02; H 5.43; N 12.33.

##### 3,3’-((2-Hydroxyphenyl)azanediyl)bis(N’-(furan-2-ylmethylene)propanehydrazide) (**15**)

White powder, yield 0.46 g (70%), m.p. 194−196 °C (from dioxane). ^1^H NMR (400 MHz, DMSO-*d_6_*) δ: 2.23−2.40 (m, 2H, CH_2_CO), 2.60−2.75 (m, 2H, CH_2_CO), 3.17−3.38 (m, 4H, 2x NCH_2_), 6.50–7.18 (m, 8H, H_Ar_); 7.69–8.07 (m, 4H, H_Ar_, N=CH); 8.95 (s, 1H, OH), 11.25, 11.27, 11.34 (3s, 2H, 2x NH); ^13^C NMR (101 MHz, DMSO-*d_6_*) δ: 30.20, 30.27 (<u>C</u>H_2_CO), 47.91, 48.12, 48.64, 48.92 (NCH_2_), 112.02, 112.10, 112.91, 112.98, 113.15, 113.22, 115.57, 115.63, 115.71, 119.16, 119.23, 119.31, 122.03, 122.53, 123.15, 124.19, 124.49, 124.86, 133.11, 133.14, 135.94, 136.03, 136.24, 136.55, 136.93, 144.74, 144.78, 144.97, 149.21, 149.40, 149.41, 152.20, 152.45, 152.77, 167.53, 167.56 (C_Ar_), 173.30, 173.32 (2x C=O). IR (KBr): ν_max_ (cm^−1^) = 3227 (OH), 3098 (2x NH), 1671 (2x C=O). Anal. Calcd. for C_22_H_23_N_5_O_5_, %: C 60.40; H 5.30; N 16.01. Found: C 60.27; H 5.17; N 15.87. HRMS m/z calculated for C_22_H_23_N_5_O_5_ [M+H]^+^: 438.1699; found: 438.1770.

##### 3,3’-((2-Hydroxyphenyl)azanediyl)bis(N’-(thiophen-2-ylmethylene)propanehydrazide) (**16**)

White powder, yield 0.49 g (70%), m.p. 212−214 °C (from dioxane). ^1^H NMR (400 MHz, DMSO-*d_6_*) δ: 2.31 and 2.68 (2t, *J* = 6.9 Hz, 4H, CH_2_CO), 3.15−3.34 (m, 4H, 2x NCH_2_), 6.67–7.63 (m, 10H, H_Ar_,H_Het_), 8.12, 8.14, 8.33 (3s, 2H 2x N=CH), 8.95 (t, *J* = 10.9 Hz, 1H, OH), 11.27, 11.29, 11.35 (3s, 2H, 2x NH); ^13^C NMR (101 MHz, DMSO-*d_6_*) δ: 30.13, 30.36, 32.36, 32.45 (<u>C</u>H_2_CO), 48.13, 48.45, 48.58, 48.93 (NCH_2_), 115.55, 115.65, 115.71, 119.14, 119.22, 119.31, 122.25, 122.65, 123.09, 124.27, 124.54, 124.81, 127.76, 127.79, 127.83, 128.11, 128.15, 128.66, 129.99, 130.05, 130.63, 130.67, 136.22, 136.45, 136.81, 138.02, 138.07, 138.99, 139.01, 139.08, 139.12, 141.20, 141.28, 152.27, 152.50, 152.73, 167.38, 167.43 (C_Ar_), 173.11, 173.13 (2x C=O). IR (KBr): ν_max_ (cm^−1^) = 3203 (OH), 3011 (2x NH), 1672 (2x C=O). Anal. Calcd. for C_22_H_23_N_5_O_3_S_2_, %: C 56.27; H 4.94; N 14.91. Found: C 52.05; H 4.74; N 14.76. HRMS m/z calculated for C_22_H_23_N_5_O_3_S_2_ [M+H]^+^: 470.1242; found: 470.1314.

##### 3,3’-((2-Hydroxyphenyl)azanediyl)bis(N’-((5-nitrothiophen-2-yl)methylene)propanehydrazide) (**17**)

Yellow powder, yield 0.6 g (72%), m.p. 221−223 °C (from methanol). ^1^H NMR (400 MHz, DMSO-*d_6_*) δ: 2.35 and 2.71 (2t, *J* = 7.2 Hz, 4H, CH_2_CO), 3.16−3.47 (m, 4H, 2x NCH_2_), 6.72–6.87 (m, 2H, H_Ar_); 6.88–6.98 (m, 1H, H_Ar_); 7.07–7.15 (m, 1H, H_Ar_); 7.33–7.50 (m, 2H, H_Ar_); 7.94–8.39 (m, 4H, H_Ar_, N=CH); 8.81, 8.86, 9.02 (3s, 1H, OH), 11.68, 11.70, 11.74, 11.76 (4s, 2H, 2x NH); ^13^C NMR (101 MHz, DMSO-*d_6_*) δ: 30.04, 30.34, 32.49, 32.65 (<u>C</u>H_2_CO), 48.11, 48.58 (NCH_2_), 115.58, 115.65, 119.08, 119.26, 122.40, 122.87, 124.69, 128.72, 128.82, 129.32, 130.44, 136.03, 136.18, 136.54, 139.49, 146.76, 146.88, 150.26, 150.63, 152.32, 152.54, 168.13 (C_Ar_), 173.61, 173.72 (2x C=O). IR (KBr): ν_max_ (cm^−1^) = 3198 (OH), 3100 (2x NH), 1672 (2x C=O). Anal. Calcd. for C_22_H_21_N_7_O_7_S_2_, %: C 47.22; H 3.78; N 17.52. Found: C 47.05; H 3.56; N 17.34.

##### 3,3’-((2-Hydroxyphenyl)azanediyl)bis(N’-(5-nitrofuran-2-ylmethylene)propanehydrazide) (**18**)

Yellow powder, yield 0.47 g (59%), m.p. 188−190 °C (from dioxane). ^1^H NMR (400 MHz, DMSO-*d_6_*) δ: 2.24−2.42 (m, 2H, CH_2_CO), 2.74 (t, *J* = 6.7 Hz, 2H, CH_2_CO), 3.20−3.41 (m, 4H, 2x NCH_2_), 6.42–7.21 (m, 6H, H_Ar_), 7.58–8.12 (m, 4H, H_Ar_ 2x N=CH), 8.84, 8.89, 9.01 (3s, 1H, OH), 11.68, 11.72, 11.76, 11.79 (4s, 2H, 2x NH); ^13^C NMR (101 MHz, DMSO-*d_6_*) δ: 30.07, 30.26, 32.53 (<u>C</u>H_2_CO), 47.85, 48.38, 48.77 (NCH_2_), 114.24, 114.35, 114.61, 114.64, 114.70, 114.97, 115.53, 115.62, 115.77, 122.18, 122.71, 123.21, 124.27, 124.59, 124.98, 130.89, 131.04, 133.84, 133.96, 136.13, 136.37, 136.75, 151.59, 151.69, 151.81, 151.85, 152.22, 152.51, 168.14, 168.20 (C_Ar_), 173.83, 173.88 (2x C=O). IR (KBr): ν_max_ (cm^−1^) = 3212 (OH), 3153 (2x NH), 1676 (2x C=O). Anal. Calcd. for C_22_H_21_N_7_O_9_, %: C 50.10; H 4.01; N 18.59. Found: C 49.94; H 3.87; N 18.35. HRMS m/z calculated for C_22_H_21_N_7_O_9_ [M+H]^+^: 528.1400; found: 528.1476.

##### 3,3’-((2-Hydroxyphenyl)azanediyl)bis(N’-(thiophen-3-ylmethylene)propanehydrazide) (**19**)

White powder, yield 0.62 g (89%), m.p. 209−211 °C (from 2-propanol). ^1^H NMR (400 MHz, DMSO-*d_6_*) δ: 2.31 and 2.72 (2t, *J* = 7.2 Hz, 4H, CH_2_CO), 3.18−3.32 (m, 4H, NCH_2_), 6.72–8.21 (m, 12H, H_Ar_, H_Het,_ 2x CH), 8.90, 8.95, 8.98 (3s, 1H, OH), 11.19, 11.28 (2s, 2H, 2x NH); ^13^C NMR (101 MHz, DMSO-*d_6_*) δ: 30.15, 30.38, 32.37, 32.54 (<u>C</u>H_2_CO), 48.26, 48.49, 48.70, 48.93 (NCH_2_), 115.53, 115.61, 115.70, 119.07, 119.18, 119.30, 122.43, 122.74, 123.07, 124.37, 124.43, 124.47, 124.58, 124.66, 124.81, 127.26, 127.30, 127.39, 127.42, 127.53, 127.86, 127.88, 136.28, 136.54, 136.98, 137.38, 137.48, 137.54, 138.64, 138.66, 141.74, 141.84, 152.40, 152.57, 152.75, 167.41, 167.49 (C_Ar_), 173.28, 173.32 (2x C=O). IR (KBr): ν_max_ (cm^−1^) = 3287 (OH), 3088 (2x NH), 1670 (2x C=O). Anal. Calcd. for C_22_H_23_N_5_O_3_S_2_, %: C 56.27; H 4.94; N 14.91. Found: C 56.05; H 4.71; N 14.84. HRMS m/z calculated for C_22_H_23_N_5_O_3_S_2_ [M+H]^+^: 470.1242; found: 470.1313.

##### 3,3’-((2-Hydroxyphenyl)azanediyl)bis(N’-(propan-2-ylidene)propanehydrazide) (**20**)

White powder, yield 0.43 g (80 %), m.p. 161−163 °C (from 2-propanol). ^1^H NMR (400 MHz, DMSO-*d_6_*) δ: 1.81, 1.82, 1.86, 1.90 (4s, 12H, 4x CH_3_); 2.32 (q, *J* = 6.9 Hz, 2H, CH_2_CO), 2.60 (t, *J* = 6.9 Hz, 2H, COCH_2_), 3.20 (q, *J* = 6.9 Hz, 4H, NCH_2_), 6.70–6.81 (m, 2H, H_Ar_), 6.85–6.95 (m, 1H, H_Ar_), 7.03–7.11 (m, 1H, H_Ar_), 8.93, 8.96, 9.01 (3s, 1H, OH), 10.02, 10.04 (2s, 2H, 2x NH); ^13^C NMR (101 MHz, DMSO-*d_6_*) δ: 17.01, 17.46, 24.96, 25.12 (4x CH_3_), 30.50, 30.64, 31.94, 32.01 (<u>C</u>H_2_CO), 47.90, 48.18, 48.84, 48.97 (NCH_2_), 115.55, 118.96, 119.07, 121.83, 122.31, 122.78, 124.14, 124.39, 124.62, 136.38, 136.63, 137.13, 150.35, 152.26, 152.61, 154.79, 154.86, 167.47 (C_Ar_), 173.51 (2x C=O). IR (KBr): ν_max_ (cm^−1^) = 3271 (OH), 3198 (2x NH), 1660 (2x C=O). Anal. Calcd. for C_18_H_27_N_5_O_3_, %: C 59.81; H 7.53; N 19.38. Found: C 59.63; H 7.36; N 19.17. HRMS m/z calculated for C_18_H_27_N_5_O_3_ [M+H]^+^: 362.2113; found: 362.2186.

##### 3,3’-((2-Hydroxyphenyl)azanediyl)bis(N’-(butan-2-ylidene)propanehydrazide) (**21**)

Light brown powder, yield 0.49 g (84 %), m.p. 81−83 °C (from 2-propanol). ^1^H NMR (400 MHz, DMSO-*d_6_*) δ: 0.82–1.11 (m, 6H, CH_2_C<u>H_3_</u>); 1.79, 1.81, 1.84, 1.88 (4s, 6H, 2x CH_3_); 2.09–2.29 (m, 4H, C<u>H_2_</u>CH_3_); 2.32 (t, *J* = 7.1 Hz, 2H, CH_2_CO), 2.63 (t, *J* = 7.1 Hz, 2H, COCH_2_), 3.21 (q, *J* = 6.9 Hz, 4H, NCH_2_), 6.67–6.82 (m, 2H, H_Ar_), 6.85–6.95 (m, 1H, H_Ar_), 7.03–7.11 (m, 1H, H_Ar_), 8.95 (1s, 1H, OH), 10.01, 10.09 (2s, 2H, 2x NH); ^13^C NMR (101 MHz, DMSO-*d_6_*) δ: 9.73, 9.78, 10.45, 10.84, 15.73, 15.88 (4x CH_3_), 22.14, 22.88, 23.31 (<u>C</u>H_2_CH_3_), 30.45, 31.43, 31.69, 31.98, 32.12 (<u>C</u>H_2_CO), 48.14, 48.50, 48.75, 48.99 (NCH_2_), 115.43, 115.54, 119.05, 122.05, 122.44, 122.74, 124.24, 124.45, 124.61, 136.56, 137.04, 152.37, 152.52, 153.73, 153.78, 158.32, 167.55 (C_Ar_), 173.72 (2x C=O). IR (KBr): ν_max_ (cm^−1^) = 3240 (OH), 3178 (2x NH), 1671 (2x C=O). Anal. Calcd. for C_20_H_31_N_5_O_3_, %: C 61.67; H 8.02; N 17.98. Found: C 61.45; H 7.88; N 17.87. HRMS m/z calculated for C_20_H_31_N_5_O_3_ [M+H]^+^: 390.2426; found: 390.2500.

##### 3,3’-((2-Hydroxyphenyl)azanediyl)bis(N’-(1-phenylethylidene)propanehydrazide) (**22**)

White powder, yield 0.61 g (84%), m.p. 187−189 °C (from 2-propanol). ^1^H NMR (400 MHz, DMSO-*d_6_*) δ: 2.09–2.27 (m, 6H, CH_3_); 2.40–2.55 (overlaps with DMSO, 2H, CH_2_CO), 2.80 (t, *J* = 7.1 Hz, 2H, COCH_2_), 3.22–3.42 (m, 4H, NCH_2_), 6.68–6.99 (m, 3H, H_Ar_), 7.07–7.18 (m, 1H, H_Ar_), 7.27–7.47 (m, 6H, H_Ar_), 7.52– 7.85 (m, 4H, H_Ar_); 8.93, 8.99 (2s, 1H, OH), 10.44, 10.45, 10.48, 10.50 (4s, 2H, 2x NH); ^13^C NMR (101 MHz, DMSO-*d_6_*) δ: 13.49, 14.03 (2x CH_3_), 30.56, 30.84, 32.18, 32.42 (<u>C</u>H_2_CO), 48.45, 48.70, 48.92 (NCH_2_), 115.47, 115.59, 119.05, 119.16, 122.53, 122.77, 124.42, 125.90, 126.26, 128.24, 128.27, 128.31, 128.87, 129.09, 136.41, 136.87, 138.11, 138.27, 147.22, 147.30, 150.97, 152.61, 168.09, 168.18 (C_Ar_), 174.29 (2x C=O). IR (KBr): ν_max_ (cm^−1^) = 3242 (OH), 3158 (2x NH), 1677 (2x C=O). Anal. Calcd. for C_28_H_31_N_5_O_3_, %: C 69.26; H 6.44; N 14.42. Found: C 69.03; H 6.21; N 14.24.

##### 3,3’-((2-Hydroxyphenyl)azanediyl)bis(N-(2,5-dimethyl-1H-pyrrol-1-yl)propanamide) (**23**)

To a solution of dihydrazide 5 (0.5 g, 1.8 mmol) in 2-propanol (25 mL), hexane-2,5-dione (0.82 g, 7.2 mmol) and a catalytic amount of acetic acid (0.1 mL) were added, and the mixture was heated under reflux for 6 h, then cooled down, and diluted with water (25 mL); the formed precipitate was filtered off, washed with water, and recrystallized from a mixture of 2-propanol and water. White powder, yield 0.53 g (67%), m.p. 200−202 °C (from 2-propanol). ^1^H NMR (400 MHz, DMSO-*d_6_*) δ: 1.95 (s, 12H, 4x CH_3_); 2.41 (t, *J* = 7.2 Hz, 4H, 2x COCH_2_), 3.34 (t, *J* = 7.2 Hz, 4H, 2x NCH_2_), 5.61 (s, 4H, 4x CH_pyr_); 6.74–6.86 (m, 2H, H_Ar_), 6.90–6.98 (m, 1H, H_Ar_), 7.09–7.16 (m, 1H, H_Ar_), 8.88 (1s, 1H, OH), 10.60 (1s, 2H, 2x NH); ^13^C NMR (101 MHz, DMSO-*d_6_*) δ: 10.93 (2x CH_3_), 31.41 (<u>C</u>H_2_CO), 47.78 (NCH_2_), 102.84, 115.68, 119.33, 123.26, 124.81, 126.70, 136.06, 152.64 (C_Ar_), 170.60 (C=O). IR (KBr): ν_max_ (cm^−1^) = 3237 (OH), 3018 (2x NH), 1672 (2x C=O). Anal. Calcd. for C_24_H_31_N_5_O_3_, %: C 65.88; H 7.14; N 16.01. Found: C 65.63; H 6.95; N 15.87. HRMS m/z calculated for C_24_H_31_N_5_O_3_ [M+H]^+^: 438.2426; found: 438.2498.

##### 5-(3-(3,5-Dimethyl-1H-pyrazol-1-yl)-3-oxopropyl)-4,5-dihydrobenzo[b][1,4]oxazepin-2(3H)-one (**24**)

To a solution of dihydrazide 5 (0.5 g, 1.8 mmol) in 2-propanol (28 mL), pentane-2,4-dione (0.9 g, 9.0 mmol) and a catalytic amount of hydrochloric acid (0.05 mL) were added, and the mixture was heated under reflux for 5 h, then cooled down. The solvent was removed under reduced pressure, the residue was poured with water (30 mL), and the formed precipitate was filtered off, washed with water and diethyl ether, and recrystallized from a mixture of 2-propanol and water. White powder, yield 0.35 g (62%), m.p. 133−135 °C (from 2-propanol). ^1^H NMR (400 MHz, DMSO-*d_6_*) δ: 2.12 and 2.44 (2s, 6H, 2x CH_3_); 2.56 and 3.22 (2t, *J* = 6.9 Hz, 4H, 2x COCH_2_), 3.38 and 3.44 (2t, *J* = 6.9 Hz, 4H, 2x NCH_2_), 6.14 (s, 1H, 4x CH_Het_); 7.00–7.33 (m, 4H, H_Ar_); ^13^C NMR (101 MHz, DMSO-*d_6_*) δ: 13.42, 14.01 (2x CH_3_), 31.62, 33.46 (<u>C</u>H_2_CO), 46.74, 54.47 (NCH_2_), 111.14, 119.45, 121.04, 123.91, 126.24, 138.16, 143.11, 146.85, 151.32, 169.94 (C_Ar_), 171.58 (C=O). IR (KBr): ν_max_ (cm^−1^) = 1748, 1729 (2x C=O). Anal. Calcd. for C_17_H_19_N_3_O_3_, %: C 65.16; H 6.11; N 13.41. Found: C 64.98; H 5.96; N 13.27. HRMS m/z calculated for C_17_H_19_N_3_O_3_ [M+Na]^+^: 336.1426; found: 336.1321.

##### 3,3’-((2-Hydroxyphenyl)azanediyl)bis(N’-((Z)-2-oxoindolin-3-ylidene)propanehydrazide) (**25**)

To a solution of hydrazide 5 (0.3, 1.06 mmol) in methanol (10 mL), isatin (0.38, 2.55 mmol) and glacial acetic acid (1 drops) were added. The reaction mixture was heated under reflux for 4 h. Precipitate was filtered off, washed with methanol, and recrystallized from 2-propanol/H_2_O mixture. Yellow powder, yield 0.48 g (83%), m.p. 156−158 °C (2-propanol/H2O mixture). ^1^H NMR (400 MHz, DMSO-*d_6_*) δ: 2.57–2.99 (m, 4H, 2x COCH_2_), 3.39 (t, *J* = 7.1 Hz, 4H, 2x NCH_2_), 6.99–7.40 (m, 10H, H_Ar_), 7.82–8.11 (m, 2H, H_Ar_), 8.94 (s, 1H, OH), 10.75 and 11.09 (2s, 4H, 4x NH); ^13^C NMR (101 MHz, DMSO-*d_6_*) δ: 31.57 (<u>C</u>H_2_CO), 48.20 (NCH_2_), 110.48, 115.28, 115.70, 119.32, 121.58, 124.36, 126.09, 126.22, 132.39, 143.60, 152.18, 164.60 (C_Ar_), 174.80, 184.90 (2x C=O). IR (KBr): ν_max_ (cm^−1^) = 3220 (OH), 3202 (4x NH), 1682, 1618 (2x C=O). Anal. Calcd. for C_28_H_25_N_7_O_5_, %: C 62.33; H 4.67; N 18.17. Found: C 62.17; H 4.42; N 17.89. HRMS m/z calculated for C_28_H_25_N_7_O_5_ [M+Na]^+^: 540.1917; found: 540.1995.

##### 5,5’-(((2-Hydroxyphenyl)azanediyl)bis(ethane-2,1-diyl))bis(1,3,4-oxadiazole-2(3H)-thione) (**26**)

A mixture of dihydrazyde 5 (0.7 g, 2.5 mmol), potassium hydroxide (2.19 g, 39 mmol), carbon disulfide (3.62 g, 47.5 mmol), and 60 mL methanol was refluxed for 24 h, and then the volatile fractions were separated under reduced pressure. The obtained residue was dissolved in water (20 mL), and the solution was acidified with acetic acid to pH 6. The formed solid was filtered off, washed with water, and recrystallized from a mixture of 2-propanol and water. White powder, yield 0.65 g (71%), m.p. 176−178 °C (from 2-propanol). ^1^H NMR (400 MHz, DMSO-*d_6_*) δ: 2.79 (t, *J* = 7.1 Hz, 4H, 2x COCH_2_), 3.40 (t, *J* = 7.1 Hz, 4H, 2x NCH_2_), 6.73 (t, *J* = 7.5 Hz, 1H, H_Ar_), 6.79 (d, *J* = 7.8 Hz, 1H, H_Ar_), 6.90 (t, *J* = 7.5 Hz, 1H, H_Ar_), 6.97 (d, *J* = 7.8 Hz, 1H, H_Ar_), 8.97 (s, 1H, OH), 14.24 (br s, 2H, 2x NH); ^13^C NMR (101 MHz, DMSO-*d_6_*) δ: 23.89 (<u>C</u>H_2_CO), 48.49 (NCH_2_), 115.96, 119.17, 123.37, 124.67, 135.16, 152.25, 162.77 (C_Ar_), 177.60 (2x C=S). IR (KBr): ν_max_ (cm^−1^) = 3193 (OH), 2966 (2x NH). Anal. Calcd. for C_14_H_15_N5O_3_S_2_, %: C 46.02; H 4.14; N 19.17. Found: C 45.87; H 3.89; N 18.96. HRMS m/z calculated for C_14_H_15_N_5_O_3_S [M+Na]^+^: 388.0616; found: 388.0508.

### Microbial strains and culture conditions

The multidrug-resistant *S. aureus* strain TCH 1516 [USA 300-HOU-MR] and pan-susceptible *S. aureus* ATCC 25923 was obtained from the American Type Culture Collection (ATCC). *A. baumannii* PKC-1027 and *E. clocea* PKC-0122 are laboratory strains of clinical origin obtained from the Institute of Infectious Diseases and Pathogenic Microbiology collection. *E. faecalis* AR-0781, *K. pneumoniae* AR-0153 and *P. aeruginosa* AR-0054 were obtained from ARisolate bank at CDC Atlanta. Prior to the experiments all microbial strains were stored in commercial cryopreservation systems at a temperature of −80 °C. The strains were cultivated on Columbia sheep blood agar (Becton Dickinson, Franklin Lakes, NJ, USA). Fungal strains were cultured on Sabouraud Dextrose agar (Becton Dickinson, Franklin Lakes, NJ, USA).

### Minimal inhibitory concentration determination

The antimicrobial activity of synthesized compounds or control antibiotics was assessed using the broth microdilution method, following the guidelines outlined by the Clinical Laboratory Standards Institute (CLSI), with modifications [29,30]. In brief, the compounds were dissolved in dimethylsulfoxide (DMSO) to attain a final concentration of 25–30 mg/mL. Dilution series were prepared in deep 96-well microplates to achieve a two-fold concentration range of 0.5, 1, 2, 4, 8, 16, 32, 64 and 128 µg/mL, utilizing cation-adjusted Mueller–Hinton broth (CAMHB) as the growth medium. For *Candida* or filamentous mold screening, dilutions were performed in RMPI/MOPS media. The microplates containing the dilution series were then inoculated with fresh cultures of each tested organism to reach a final concentration of 5 × 10^4^ CFU (colony-forming units) of the test organism in media containing 1% DMSO and 1× drug concentration, with a volume of 200 µL per well. Wells that were inoculated with media containing 1% DMSO served as positive controls. Subsequently, the microplates were incubated at 35 ± 1 °C for 18 ± 2 h. Following the incubation period, the plates were examined using a manual microplate viewer (Sensititre Manual Viewbox, United States). The minimal inhibitory concentration (MIC) was defined as the lowest concentration (µg/mL) of the tested drug that completely inhibited the growth of the test organism. All experiments were conducted in duplicate with three technical replicates for each condition.

## Results

### Synthesis of N-Substituted β-Amino Acid Derivatives

In the first stage of this work, by using a well-known methodology described in the publication [31], the initial compound **5** were prepared. According to the methodology, the reaction of 2-aminophenol (**1**) with acrylic acid in water at reflux afforded intermediates 3,3′-((2-hydroxyphenyl)azanediyl)di(propanoic)acid (**2**) (Scheme 1).

**Scheme 1.**
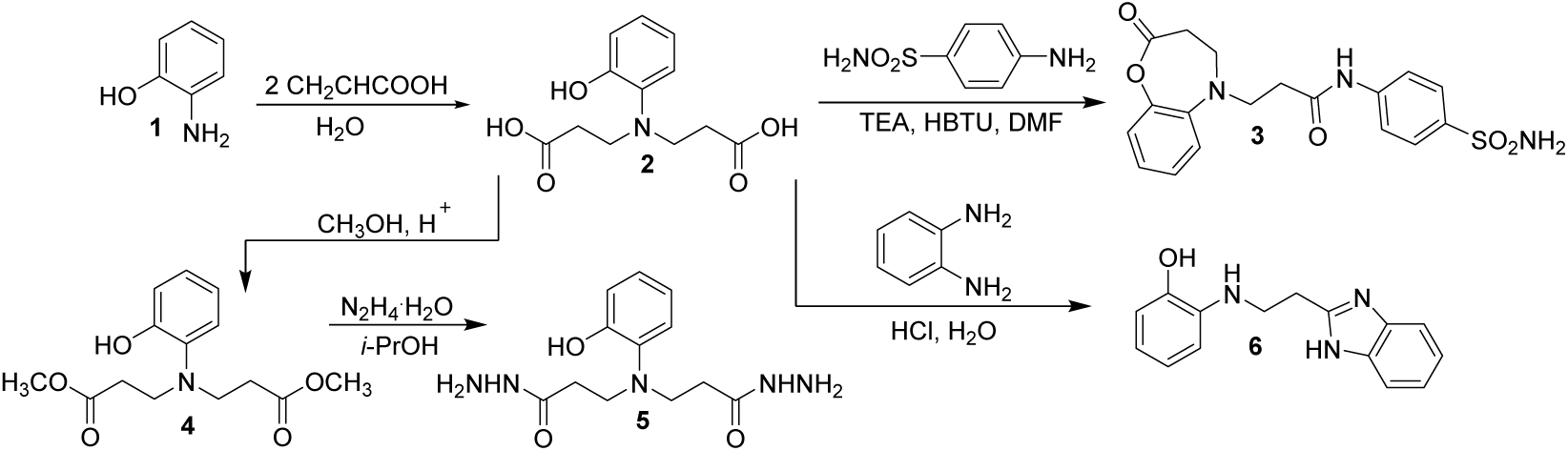
Synthesis of compounds 2–6.

The attempt to synthetize the diamide **3** led to unexpected product. However, the diacid **2** cyclizes under these reaction conditions to the oxadiazepine moiety bearing derivative **3**. Compound **3** was prepared by direct coupling of acids **2** with the sulphanilamide using HBTU as the coupling reagent and triethylamine as the base. The reaction were performed in dimethylformamide at room temperature. The product **3** was isolated by the dilution of the reaction mixture with water and were characterised using NMR, HRMS, IR spectroscopy and elemental analysis. Comparison of the ^1^H NMR spectra of product **3** with the compound **2** has revealed, that proton singlet of hydroxy group at 8.77 ppm in the spectra of diacid **2** have been dissapered with the formation of the oxazepine fragment. The structure of the obtained compound **3** is also confirmed by the results of the HRMS m/z calculated for C_18_H_19_N_3_O_5_S [M+H]^+^: 390.1045; found: 390.1119.

In continuation of our interest in the chemistry of *N*-substituted *β*-amino acids, dimethyl ester **4** was synthesized through esterification of 3,3′-((2-hydroxyphenyl)azanediyl)di(propanoic)acid (**2**) with an excess of methanol in the presence of a catalytic amount of sulfuric acid. Dihydrazide **5** was obtained through hydrazinolysis of dimethyl ester **4** in propan-2-ol under reflux.

Condensation of dihydrazide **5** with aromatic aldehydes and ketones gave the corresponding hydrazones **7**–**22.** The structures of hydrazones **7**–**22** have been established mainly on the basis of ^1^H and ^13^C NMR spectra (Scheme 2).

**Scheme 2.**
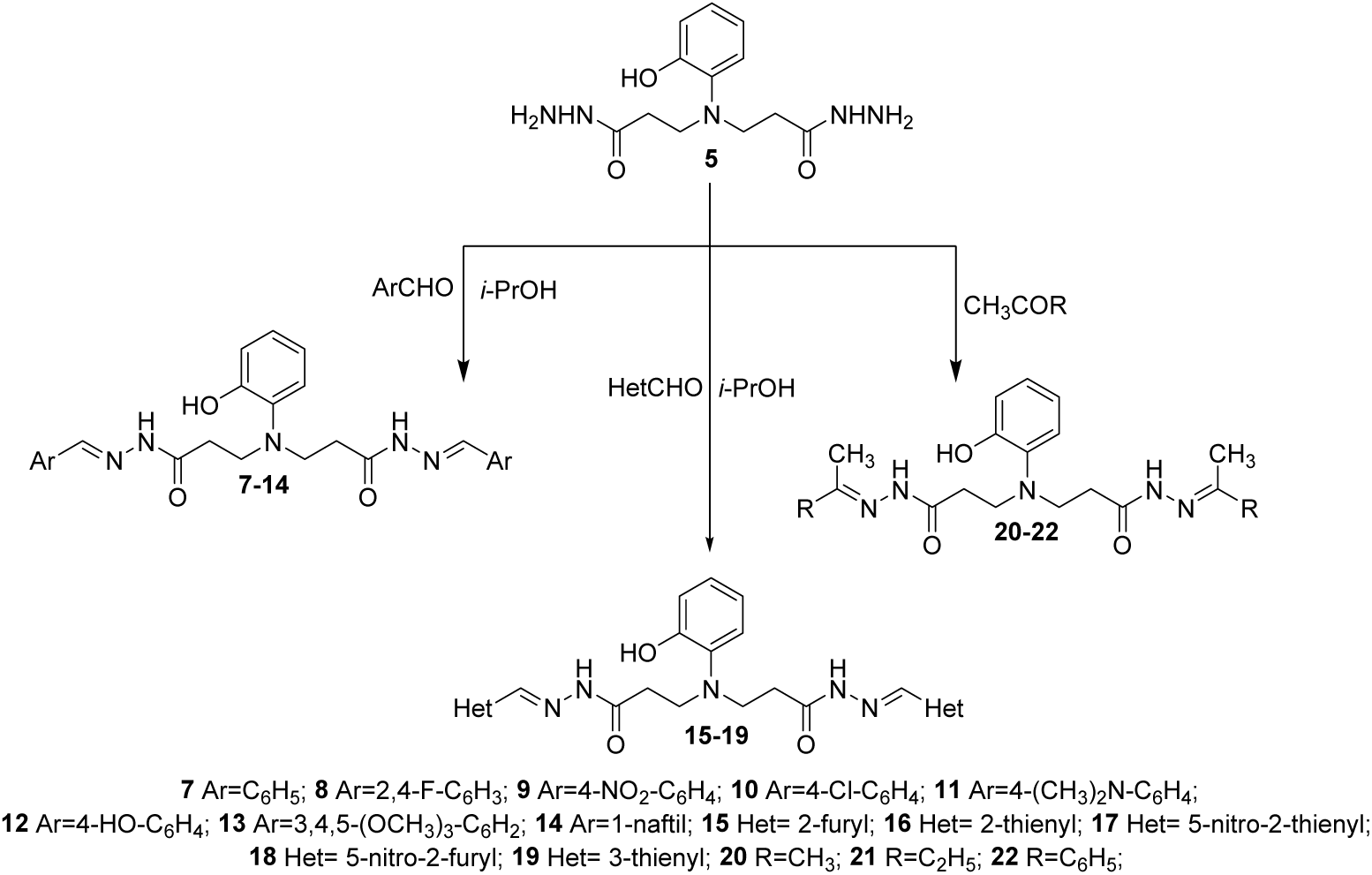
Synthesis of compounds 7–22.

In the next stage of this work, condensation reactions of dihydrazide **5** with various dicarbonyl compounds and carbon disulfite were performed. The reaction of dihydrazide **5** with acetylacetone, isatin and carbon disulfide resulted in the formation of compounds **23, 25** and **26** of the expected structure, each containing two identical heterocyclic fragments, while in the reaction with 2,5-hexadione, cyclization also took place with the participation of the hydroxy group in the *o*-position, forming the corresponding 5-(3-(3,5-dimethyl-1*H*-pyrazol-1-yl)-3-oxopropyl)-4,5-dihydrobenzo[*b*][1,4]oxazepin-2(3*H*)-one **(24)** (Scheme 3).

**Scheme 3.**
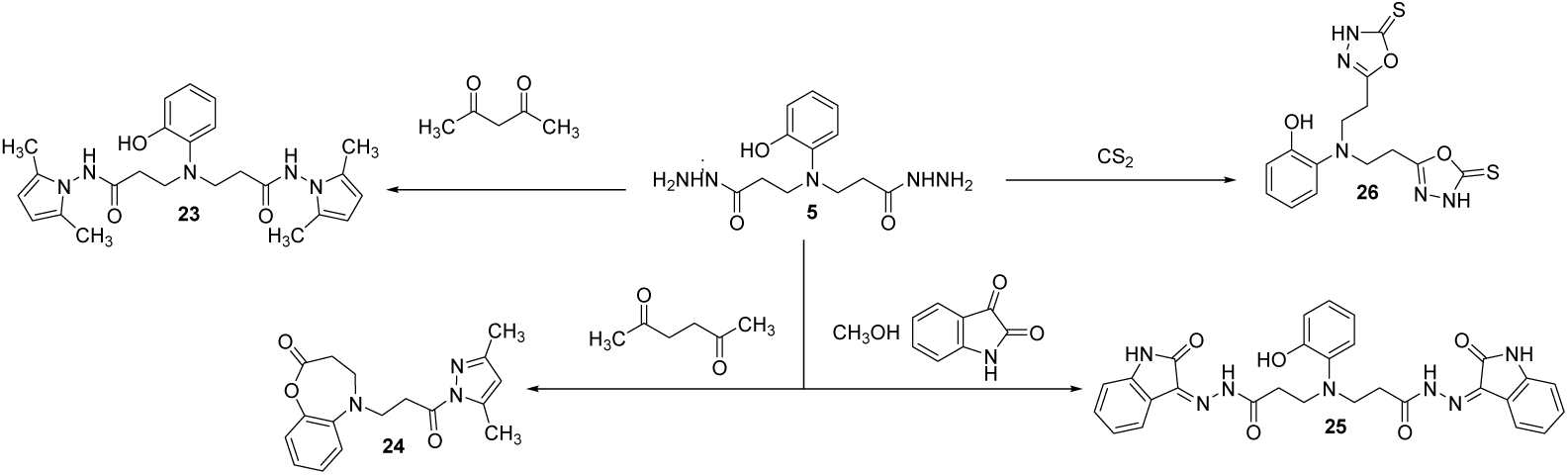
Synthesis of compounds 23–26.

Comparison of the ^1^H NMR spectra of product **24** with the compound **5** has revealed, that proton singlet of hydroxy group at 9.01 ppm in the spectra of dihydrazide **5** have been dissapered with the formation of the oxazepine fragment. The structure of the obtained compound **24** is also confirmed by the results of the HRMS - m/z calculated for C_17_H_19_N_3_O_3_ [M+Na]^+^: 336.1426; found: 336.1321.

### 2-Hydroxyphenyl propanoic acid derivatives shows Gram-positive bacteria-directed activity

After successfully synthesizing and characterizing a series of 2-hydroxyphenyl propanoic acid derivatives (compounds **2–26**), we evaluated their *in vitro* antimicrobial activity. The compounds were screened using a laboratory strain collection of ESKAPE pathogens, including *Enterococcus spp., Staphylococcus aureus, Klebsiella pneumoniae, Acinetobacter baumannii, Pseudomonas aeruginosa,* and *Enterobacter spp* and their minimal inhibitory concentration (MIC) was determined. These pathogens were selected due to their clinical significance and the presence of genetically defined and emerging antimicrobial resistance mechanisms, while the control antibiotics (vancomycin, gentamycin, meropenem, and cefazolin) were selected to represent the antibacterial agents used in the clinical setting to treat infections caused by Gram-negative and Gram-positive agents.

Starting compound **2** demonstrated no antimicrobial activity against tested bacterial and fungal strains (Table 1, Table S1). Oxadiazepine derivative **3** demonstrated moderate antimicrobial activity against *Enterococcus faecalis* AR-0781 and *Staphylococcus aureus* TCH-1516, with a minimum inhibitory concentration (MIC) of 64 µg/mL. Dimethyl ester **4** showed no antimicrobial activity against any of the tested bacterial and fungal strains (Table 1, Table S1). The dihydrazide **5** exhibited activity against *S. aureus* TCH-1516 (MIC 64 µg/mL) but not against *E. faecalis* AR-0781 or any tested Gram-negative pathogens. Benzimidazole **6** demonstrated moderate activity against *E. faecalis* AR-0781 (MIC 64 µg/mL) and weak activity against *S. aureus* TCH-1516 (MIC 128 µg/mL).

**Table 1.** The *in vitro* antimicrobial activity of *N*-substituted *β*-amino acid derivatives **2–26** against panel of multidrug-resistant bacterial isolates.

| Compound | Minimal Inhibitory Concentration ($\mu\text{g/mL}$) | | | | | |
| --- | --- | --- | --- | --- | --- | --- |
|  | *E. faecalis*<br>AR-0781<sup>A</sup> | *S. aureus*<br>TCH-1516<sup>B</sup> | *K. pneumoniae*<br>AR-0153<sup>C</sup> | *A. baumannii*<br>PKC-1027<sup>D</sup> | *P. aeruginosa*<br>AR-0054<sup>E</sup> | *E. cloacae*<br>PKC-0122<sup>F</sup> |
| **2** | 128> | 128> | 128> | 128> | 128> | 128> |
| **3** | 64 | 64 | 128> | 128> | 128> | 128> |
| **4** | 128> | 128> | 128> | 128> | 128> | 128> |
| **5** | 128> | 64 | 128> | 128> | 128> | 128> |
| **6** | 64 | 128 | 128> | 128> | 128> | 128> |
| **7** | 128> | 128> | 128> | 128> | 128> | 128> |
| **8** | 128> | 32 | 128> | 128> | 128> | 128 |
| **9** | 16 | 8 | 128> | 128> | 128> | 128> |
| **10** | 128> | 128> | 128> | 128> | 128> | 128> |
| **11** | 128> | 128> | 128> | 128> | 128> | 128> |
| **12** | 128 | 16 | 128> | 128> | 128> | 128> |
| **13** | 128> | 128> | 128> | 128> | 128> | 128> |
| **14** | 128> | 8 | 128> | 128> | 128> | 128> |
| **15** | 128> | 128> | 128> | 128> | 128> | 128> |
| **16** | 128> | 64 | 128> | 128> | 128> | 128> |
| **17** | 128> | 8 | 128> | 128> | 128> | 128> |
| **18** | 128> | 16 | 128> | 128> | 128> | 128> |
| **19** | 128> | 128> | 128> | 128> | 128> | 128> |
| **20** | 128> | 128> | 128> | 128> | 128> | 128> |
| **21** | 128> | 128> | 128> | 128> | 128> | 128> |
| **22** | 128> | 128> | 128> | 128> | 128> | 128> |
| **23** | 128> | 128> | 128> | 128> | 128> | 128> |
| **24** | 128 | 32 | 128> | 128> | 128> | 128> |
| **25** | 64 | 32 | 128> | 128> | 128> | 128> |
| **26** | 8 | 4 | 128> | 128> | 128> | 128> |
| **Vancomycin** | 64> | 2 | N/A | N/A | N/A | N/A |
| **Gentamycin** | 16 | 32 | 64 | 128> | 128 | 64 |
| **Meropenem** | 2 | 2 | 64 | 64 | 64 | 2 |
| **Cefazolin** | 8 | 32 | 128> | 128> | 128> | 32 |

A. Vancomycin-resistant *E. faecalis* harboring *tet(L)*, *tet(M)*, *VanA* resistance genes.
B. Meticillin-Resistant *S. aureus* USA300 lineage harboring *mecA* and Panton-Valentine leucocidin *pvl*.
C. Carbapenem-resistant *K. pneumoniae* harboring *aac(3)-IId*, *aadA2*, *armA*, *cmlA1*, *CTX-M-15*, *dfrA1*, *dfrA12*, *dfrA14*, *fosA*, *mph(E)*, *msr(E)*, *NDM-1*, *oqxA*, *OXA-1*, *OXA-232*, *OXA-9*, *strA*, *strB*, *sul1*, *sul2*, *TEM-1A* resistance genes.
D. Colistin-resistant *A. baumannii* complex blood isolate.
E. Carbapenem-resistant *P. aeruginosa* harboring *ant(2'')-Ia*, *aph(3')-IId*, *aph(6)-Id*, *bcr1*, *bcr1*, *bcr1*, *catB7*, *fosA*, *mexA*, *mexE*, *mexX*, *OXA-396*, *PDC-3*, *sul1*, *sul1*, *sul1*, *tet(A)*, *tet(R)*, *tet(R)*, *tet(R)*, *VIM-4* resistance genes.
F. Extended-spectrum beta-lactamases (ESBL) producing blood isolate of *E. cloacea* harboring *CTX-M-15*, *aac(3)-IId*, *aadA2* resistance genes.

To characterize the *in vitro* structure-dependent effects of aromatic substitutions on antimicrobial activity, hydrazide **5** was used as a starting compound. The incorporation of an aryl substituent resulted in compound **7**, which showed no antimicrobial activity against all tested bacterial and fungal isolates (MIC >128 µg/mL) (Table 1, Table S1). The incorporation of a 2,4-difluorophenyl substituent (compound **8**) demonstrated activity against methicillin-resistant *S. aureus* TCH-1516 (MIC 32 µg/mL) (MRSA) but not vancomycin-resistant *E. faecalis* AR-0781 (MIC >128 µg/mL) (VRE). Interestingly, compound **8** showed weak activity against the ESBL-producing *E. cloacae* PKC-0122 strain (MIC 128 µg/mL). In addition, the incorporation of a 4-nitrophenyl substituent (compound **9**) greatly enhanced the in vitro antimicrobial activity against *E. faecalis* AR-0781 (MIC 16 µg/mL) and *S. aureus* TCH-1516 (MIC 8 µg/mL), although it resulted in a loss of activity against *E. cloacae* PKC-0122 (MIC >128 µg/mL). Furthermore, the addition of a 4-chlorophenyl (compound **10**) or 4-(dimethylamino)phenyl (compound **11**) substituent resulted in a loss of antimicrobial activity against all tested isolates (MIC >128 µg/mL). The incorporation of a 4-hydroxyphenyl substituent (compound **12**) showed weak activity against *E. faecalis* AR-0781 (MIC 128 µg/mL) and favorable activity against *S. aureus* TCH-1516 (MIC 16 µg/mL). The 3,4,5-trimethoxyphenyl derivative (compound **13**) showed no antimicrobial activity, while the addition of a 1-naphthyl substituent (compound **14**) resulted in favorable activity against *S. aureus* TCH-1516 (MIC 8 µg/mL) (Table 1).

To better understand the function of aromatic substitutions on antimicrobial activity, we further generated a series of compounds with heterocyclic substituents (compounds **15**–**19**). Interestingly, compounds bearing heterocyclic substitutions demonstrated activity only against the *S. aureus* TCH-1516 strain. Compound **15**, bearing a 2-furyl substituent, showed no antimicrobial activity (MIC >128 µg/mL). The incorporation of a 2-thienyl substituent resulted in compound **16** with weak antimicrobial activity against the *S. aureus* TCH-1516 strain (MIC 64 µg/mL).

The incorporation of a 4-nitro group on the 2-thienyl (compound **17**) enhanced antimicrobial activity against *S. aureus* TCH-1516 (MIC 8 µg/mL), while the replacement of the 4-nitro-2-thienyl with a 5-nitro-2-furyl (compound **18**) reduced activity against *S. aureus* TCH-1516 (MIC 16 µg/mL). Surprisingly, the addition of a 3-thienyl substituent (compound **19**) diminished antimicrobial activity against *S. aureus* TCH-1516 (MIC >128 µg/mL), demonstrating that the 2-thienyl position is crucial for *S. aureus* TCH-1516-directed activity. Finally, the methyl group adjacent substitutions resulted in compounds **20–22**, with no antimicrobial activity against the tested strains (Table 1).

To characterize the effect of heterocyclic substituents on the *in vitro* antimicrobial activity, we generated a series of compounds bearing known heterocyclic pharmacophores such as dimethylpyrole, isatin or thiosemicarbazide. Compound **23** bearing two identical dimethylpyrole substitutions demonstrated no antimicrobial activity against tested bacterial strains (MIC 128> µg/mL). Oxadiazepine derivative **24** bearing dimethyl pyrazole substitution demonstrated weak activity against *E. faecalis* AR-0781(MIC 128 µg/mL) and favorable activity against *S. aureus* TCH-1516 (MIC 32 µg/mL). Compound **25** bearing isatin substituent showed activity against both *E. faecalis* AR-0781(MIC 64 µg/mL) and *S. aureus* TCH-1516 (MIC 32 µg/mL). Thiosemicarbazide derivative **26** showed potent antimicrobial activity against *E. faecalis* AR-0781(MIC 8 µg/mL) and *S. aureus* TCH-1516 (MIC 4 µg/mL) that was comparable to control antibiotics.

We then generated a series of compounds bearing known heterocyclic pharmacophores such as dimethylpyrrole, isatin, or thiosemicarbazide. Compound **23**, bearing two identical dimethylpyrrole substitutions, demonstrated no antimicrobial activity against the tested bacterial strains (MIC >128 µg/mL). Oxadiazepine derivative **24**, bearing a dimethylpyrazole substitution, demonstrated weak activity against *E. faecalis* AR-0781 (MIC 128 µg/mL) and favorable activity against *S. aureus* TCH-1516 (MIC 32 µg/mL).

Compound **25**, bearing an isatin substituent, showed activity against both *E. faecalis* AR-0781 (MIC 64 µg/mL) and *S. aureus* TCH-1516 (MIC 32 µg/mL). The thiosemicarbazide derivative **26** showed potent antimicrobial activity against *E. faecalis* AR-0781 (MIC 8 µg/mL) and *S. aureus* TCH-1516 (MIC 4 µg/mL), which was similar to Gram-positive bacteria-targeting antibiotics used for this study.

These results demonstrates that *N*-substituted *β*-amino acid derivative **26** could be further explored as starting compound to generate a sub-library of compounds potentially targeting multidrug-resistant Gram-positive pathogens with emerging antimicrobial resistance mechanisms.

## Discussion

In this study we describe the synthesis of novel *N*-substituted *β*-amino acid derivatives bearing 2-hydroxyphenyl moieties as promising antimicrobial candidates targeting drug-resistant Gram-positive priority pathogens with genetically defined resistance mechanisms. This study identifies a promising *N*-substituted *β*-amino acid-based scaffold with broad Gram-positive pathogens-directed activity for further hit to lead optimization.

The ESKAPE group pathogens, comprising *Enterococcus* species*., Staphylococcus aureus, Klebsiella pneumoniae*, *Acinetobacter baumannii*, *Pseudomonas aeruginosa*, and *Enterobacter* species, are extremely challenging for their role in hospital-acquired infections and multidrug resistance [2,4, 32]. Among these, vancomycin-resistant *Enterococcus* (VRE) and methicillin-resistant *Staphylococcus aureus* (MRSA) pose significant clinical challenges due to their extensive resistance to multiple antibiotics, complicating therapeutic strategies and leading to fatal outcomes [10,13,30]. Epidemiologically, VRE and MRSA are highly prevalent in nosocomial environments, contributing to severe infections, increased morbidity and mortality rates, and substantial healthcare expenditures [30]. The persistent difficulties in managing infections caused by these pathogens highlight the critical need for the development of novel antimicrobial agents.

*N*-substituted *β*-amino acid derivatives bearing phenolic groups represent a promising class of compounds due to their potential as pharmacophores [26,27, 31]. These derivatives exhibit strong hydrogen-bonding capabilities and the ability to potentially form stable interactions with microbial cellular targets, thereby leading to the antimicrobial activity [26,27, 31]. The incorporation of phenolic groups confers additional bactericidal properties, while aromatic or heterocyclic substitutions can further augment the binding affinity and specificity of these molecules [26, 28, 30]. In this study, we successfully synthesized a series of *N*-substituted *β*-amino acid derivatives bearing a 2-hydroxyphenyl group. These compounds were designed to represent a range of aromatic and heterocyclic substituents and exhibited a structure-dependent antimicrobial activity, predominantly against multidrug-resistant Gram-positive pathogens. Among the synthesized compounds, compounds **9** (R = 4-nitrophenyl), **17** (R = 5-nitro-2-thienyl), **18** (R = 5-nitro-2-furyl), thiosemicarbazide **16**, and **26** demonstrated the most potent activity against *S. aureus* MRSA USA300 strain TCH-1516, with minimum inhibitory concentration values ranging from 4 to 16 µg/mL. The enhanced antimicrobial activity observed for these compounds can be attributed to the physicochemical properties of the substituents as well as geenral *N*-substituted *β*-amino acid or 2-hydroxyphenyl pharmacophores. The nitro groups present in compounds **9, 17**, and **18** are strong electron-withdrawing moieties, which increase the electrophilicity of the aromatic ring and may enhance the binding affinity to various bacterial target sites. Furthermore, the heterocyclic rings in compounds **17** (thienyl) and **18** (furyl) provide additional molecular properties for interaction through π-π stacking and hydrogen bonding with bacterial enzymes or membrane components. Compound **26**, bearing a thiosemicarbazide group, demonstrated significant antimicrobial activity against vancomycin-resistant *E. faecalis* AR-0781, with an activity profile comparable to that of control antibiotics.

The thiosemicarbazide moiety, containing sulfur and nitrogen atoms, is potentially involved in the formation of strong intermolecular interactions, including hydrogen bonding and coordination with metal ions, which may contribute to its potent bioactivity. In our previous studies, we successfully synthesized *N*-substituted *β*-amino acid derivatives bearing a 3-hydroxyphenyl core, which exhibited notable antimicrobial activity against Gram-positive drug-resistant pathogens as well as drug-resistant *Candida* species [28]. These findings suggest that the *N*-substituted *β*-amino acid scaffold represents a promising platform for the discovery of novel antimicrobial agents. Importantly, the position of the hydroxyl group on the phenolic ring appears to play a critical role in modulating the antimicrobial spectrum [34, 35]. Specifically, phenolic groups with hydroxyl substitutions at different positions on the aromatic ring may influence both antibacterial and antifungal activities. Substitutions in the meta-position, as seen in our previous derivatives, seem to enhance activity against Gram-positive bacteria, while the incorporation of hydroxyl groups at other positions could potentially broaden the spectrum to include fungal pathogens. The data presented in this study underscore the significant potential of *N*-substituted *β*-amino acid derivatives as novel antimicrobial agents, particularly due to their selective activity against multidrug-resistant Gram-positive pathogens. The structure-activity relationship observed, especially with regard to the position of the phenolic hydroxyl group and the nature of the substituents, highlights the critical role of molecular design in optimizing antimicrobial efficacy. The strong activity exhibited by compounds containing nitro-aromatic and heterocyclic moieties reinforces the importance of electron-withdrawing groups and π-stacking interactions in effectively targeting resistant Gram-positive bacterial strains. The broad-spectrum activity of compound **26** against both vancomycin-resistant *Enterococcus faecalis* and methicillin-resistant *Staphylococcus aureus* suggests that this scaffold could serve as a foundation for further optimization to address other high-priority pathogens, including those identified by the World Health Organization as critical threats. However, despite the promising results, this study has several limitations. First, additional screening is needed against a broader range of bacterial strains with diverse drug-resistant phenotypes to fully assess the efficacy of the most promising compounds. Second, although strong activity was observed against *S. aureus* and *E. faecalis*, there remains a gap in our understanding of how these compounds perform against other clinically significant Gram-positive pathogens, such as *Streptococcus* species or other Staphylococcus strains.

Finally, further studies are required to elucidate the specific bacterial targets of these *N*-substituted *β*-amino acid derivatives, which will be crucial for further advancing their development as therapeutic candidates for the early pre-clinical screening.

## Conclusions

Collectively, in this study, we synthesized and evaluated novel 3,3’-((2-hydroxyphenyl)azanediyl)dipropionic acid derivatives containing ester, hydrazine, hydrazones, benzimidazole, dimethylpyrrole, dimethylpyrazole and oxadezipine moieties, which exhibited significant antimicrobial activity against multidrug-resistant Gram-positive pathogens, particularly MRSA and VRE. The structure-activity relationship demonstrated the importance of substituent selection, with electron-withdrawing nitro-aromatic and heterocyclic groups enhancing antimicrobial activity.

The broad-spectrum activity of compound **26**, underscores the potential of this scaffold for further optimization. While the results are promising, further screening across additional drug-resistant resistant Gram-positive isolates and molecular studies to elucidate the precise molecular targets are needed to fully enhance the antimicrobial potential of these compounds. These findings suggest that *N*-substituted *β*-amino acid derivatives represent a promising platform for the development of new antimicrobial candidates.

## Acknowledgement

We are thankful for the supportive staff of Weill Cornell Medicine of Cornell University and Kaunas University of Technology for their immense technical help and support during this study.

## Supplementary Materials

Figure S1–S49. ^1^H and ^13^C NMR spectra of compounds **2−26** (in DMSO-*d_6_*). Table S1. The *in vitro* antimicrobial activity of *N*-substituted *β*-amino acid derivatives **2–26** against panel of fungal pathogens.

## Notes

### Competing Interest Statement

The authors have declared no competing interest.

